# Boosting Gene Expression Clustering with System-Wide Biological Information: A Robust Autoencoder Approach

**DOI:** 10.1101/214122

**Authors:** Hongzhu Cui, Chong Zhou, Xinyu Dai, Yuting Liang, Randy Paffenroth, Dmitry Korkin

**Affiliations:** Bioinformatics and Computational Biology Program, Worcester Polytechnic Institute, Worcester, MA, USA 010609; Data Science Program, Worcester Polytechnic Institute, Worcester, MA, USA 010609; Mathematics Department, Worcester Polytechnic Institute, Worcester, MA, USA 010609; Computer Science Department, Worcester Polytechnic Institute, Worcester, MA, USA 010609

## Abstract

Gene expression analysis provides genome-wide insights into the transcriptional activity of a cell. One of the first computational steps in exploration and analysis of the gene expression data is clustering. With a number of standard clustering methods routinely used, most of the methods do not take prior biological information into account. In this paper, we propose a new approach for gene expression clustering analysis. The approach benefits from a new deep learning architecture, Robust Autoencoder, which provides a more accurate high-level representation of the feature sets, and from incorporating prior biological information into the clustering process. We tested our approach on two distinct gene expression datasets and compared the performance with two widely used clustering methods, hierarchical clustering and k-means, as well as with a recent deep learning clustering approach. As a result, our approach outperformed all other clustering methods on the labeled yeast gene expression dataset. Furthermore we showed that it is better in identifying the functionally common clusters than k-means on the unlabeled human gene expression dataset. The results demonstrate that our new deep learning architecture could generalize well the specific properties of gene expression profiles. Furthermore, the results confirm our hypothesis that the prior biological network knowledge could be helpful in the gene expression clustering task.

## Introduction

Gene expression quantification and analysis using DNA microarrays, RNA sequencing (RNA-Seq), and other methods (Edfors, et al., 2016; Lockhart and Winzeler, 2000; Wang, et al., 2009) have been proved to be an exceptionally powerful tool to quantitatively study the relationships among sets of genes. Global gene expression analysis provides quantitative information about the protein and mRNA abundance across the whole organism and in the individual tissues and cells (Lovén, et al., 2012), allowing to explore a wide range of biological processes (Belacel, et al., 2006). Capturing the gene expression patterns can help studying molecular mechanisms implicated in diseases and cellular responses to drug treatment, thus facilitating drug discovery and development (Lovén, et al., 2012). Global analysis of the gene expression data has been carried out by a number of supervised and unsupervised machine learning methods (Kuo, et al., 2004; Lyons-Weiler, et al., 2003). An intuitive approach to analysis of the massive volumes of expression data is to first group the genes into smaller subsets based on common expression patterns they share, and without any preliminary knowledge of what each of these groups should include. Unsupervised learning, or clustering, methods are well-suited to address this problem (D'Haeseleer, 2005).

Until recent, clustering of the genes expression data has been commonly carried out using the classical unsupervised learning methods, such as k-means or Expectation Maximization (EM) algorithms (Hartigan and Wong, 1979; Moon, 1996). At the same time, deep learning has made great strides in advancing both supervised and unsupervised learning, becoming routine methods in image recognition (Ciregan, et al., 2012), natural language processing (Collobert and Weston, 2008), and most recently in bioinformatics and genomics (Chen, et al., 2016; LeCun, et al., 2015). Autoencoder is one of the commonly used deep architectures, and it has been proven successful to learn low-dimensional representations of biological data (Chen, et al., 2016). However, an autoencoder is insensitive to the outliers, which are widely present in the gene expression data. As a result, such architecture might not generalize well the specific properties of gene expression data. Furthermore, most of the current clustering methods do not take into account the prior biological information that could guide the clustering procedure.

In the past decade, substantial improvements have been made in utilizing high-throughput ‘‘-omics’’ to map most components of cellular networks (Barabasi and Oltvai, 2004; Cui, et al., 2015). Among them, human protein interactome and its edgotyping studies have attracted major attention (Rolland, et al., 2014). Network properties of the interactome have provided insights into the system-wide biological properties and the interactome evolution (Alhindi, et al., 2017; Han, et al., 2004). Of special interest is a property that is also found in many real-world networks, the community structure (Leskovec, et al., 2008), in which the network nodes are joined together in tightly knit groups, while the groups themselves are only loosely connected with each other. One of the key ideas behind our work is incorporating the gene community information for the tested gene sets into the clustering process; we expect that such information would improve the clustering accuracy.

Here, we propose a novel protocol, which combines a new deep architecture with the prior biological knowledge for gene expression clustering analysis. Our protocol could be divided into two main stages. First, we use a deep network to learn important characteristics of the gene expression profiles. We leverage a new autoencoder method, Robust autoencoder (Zhou, 2017). The approach is designed to extract more robust features from the input data. Once the network is trained, the low dimensional representation of the gene expression profile is used for the clustering task. In the second stage, we define a network-based metric which allows introducing the community information of each gene in the network into our clustering process. The hypothesis behind this idea is that, if two genes are in the same network community, then they are more likely to communicate with each other and share the same expression pattern. Our new clustering protocol is based on the Eisen clustering, an established hierarchical clustering approach (Eisen, et al., 1998).

We evaluated our method on two distinct gene expression datasets, one with external labels and the other one unlabeled. Specifically, we compared the performance of our method for gene expression clustering with two traditional clustering methods that are commonly used for the gene expression analysis, k-means and hierarchical clustering. We found that our method outperformed the traditional clustering methods on both labeled and unlabeled datasets. Furthermore, the proposed approach was more accurate than a deep learning autoencoder method. The results demonstrate that the new deep architecture could capture the high-level features from the gene expression profiles. Furthermore, the results confirm our hypothesis that the prior biological network knowledge could be utilized for optimizing the gene expression clustering task.

## Methods

### Problem Formulation

The clustering tasks in our approach (Fig. 1) are carried-out using unsupervised learning methods. For a given similarity measure defined in an unsupervised learning method, the objects belonging to the same cluster are more similar to each other than to those ones from other clusters. In the case of gene expression data clustering, a cluster may contain a number of genes or samples with similar expression patterns. After the preprocessing stage, the data are presented as a matrix *X*={*x*_*ij*_}, where *x*_*ij*_ stands for an expression level of gene *i* from sample *j* at a specific time point or in a specific condition. The clustering of gene expression data can be divided into two main categories: gene-based clustering and sample-based clustering (Jiang, et al., 2004). In this work, we focus on the gene-based clustering. The goal is to group genes with similar expression patterns (co-expressed genes). The expression patterns, in turn, will be used to help in our understanding of gene function, gene regulation, and cellular processes.

**Figure 1:**
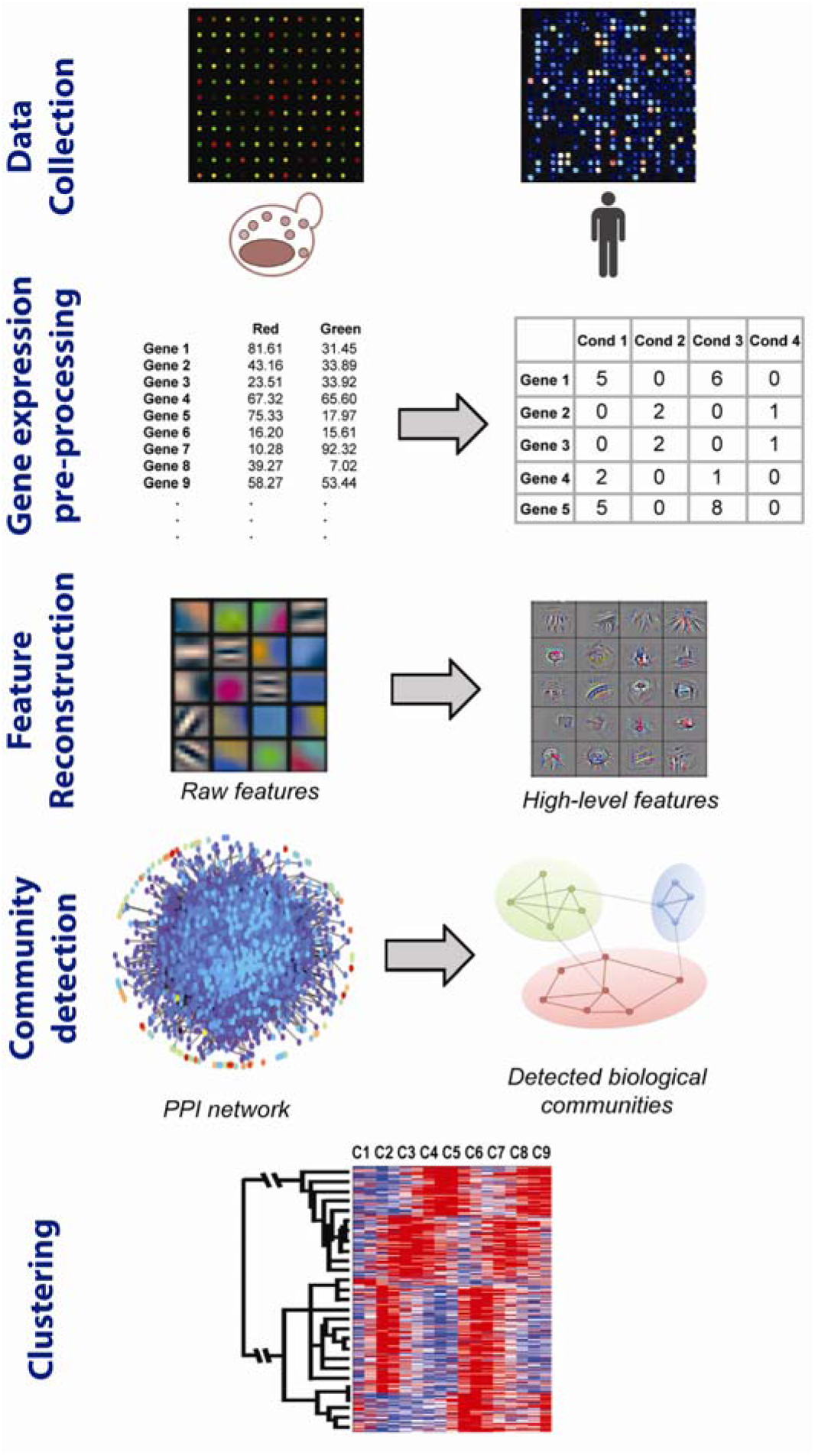
General workflow of our protocol after acquiring the raw gene expression data. Our method consists of four basic steps: input data pre-processing, feature reconstruction using deep architectures, detecting community structure from the network, and incorporating gene network community information into clustering. The two datasets used in this study are gene expression dataset for the yeast cell cycle and human gene expression data from the genomics of drug sensitivity in cancer study.

### Gene Expression Datasets

To test our approach on the real-world data, we used two distinct large-scale datasets. The first dataset includes gene expression for the yeast cell cycle (Yeung and Ruzzo, 2001). It is organized in 17 time stamps for a set of 420 genes in yeast. Based on the gene functional categories, each gene was assigned to one or more "phases". We removed the gene expression profiles for the genes that were assigned to more than one phase, resulting in a subset of 384 genes that were partitioned into 5 phases of cell cycle. The yeast dataset is widely used in practice to assess the clustering quality using the five phases assignment as an external criterion (Gupta, et al., 2015; Jiang, et al., 2004). The second dataset is obtained from the Genomics of Drug Sensitivity in Cancer (GDSC) study (Yang, et al., 2013). The dataset captures the gene expression profiles of different human cancer cell lines in response to drug compounds. It consists of 17,419 genes expressed in 83 cell lines. Overall, these two datasets differ in several principal aspects. First, the datasets are of substantially different sizes. In addition, the first dataset is time series data, while the second dataset is from different cancer cell lines. Finally, the datasets come from two different species.

### Construction of the PPI networks

To extract the community information for the gene set and link it to the expression data, we studied the protein products of these genes in the context of the physical protein-protein interaction (PPI) network (Fig. 1). To this end, two PPI networks are used: HINT yeast network (Das and Yu, 2012) and the human interactome project network (HI-II-14) (Rolland, et al., 2014). HINT network is organized as a database of high-quality protein-protein interactions collected from several databases manually as well as using an automated protocol. The comprehensive coverage of the interactome makes it possible to fully understand the network properties of the yeast genes. The human interactome HI-II-14 is another recently released source of PPI data. It is constructed through mapping binary PPIs obtained by systematically interrogating all pairwise combinations of human proteins using yeast two-hybrid high-throughput experiments. For each network, we run the community detection algorithm and apply the extracted community information during clustering.

### Community Detection Algorithms and Weighting Strategy

Community structure could be viewed as a subnetwork of nodes that are more densely connected compared to the parts of the network (Fig. 2). It is a common characteristic in many physical networks, including the Internet and World Wide Web, social networks, and different kinds of biomolecular networks (Leskovec, et al., 2008). Determining these community structures in a network can provide insight into the structural and functional organization of the network and can be useful in improving graph algorithms, such as spectral clustering (Leskovec, et al., 2008). In a basic community detection setting, a network node is defined as belonging to at most one community. The majority of community detection methods adopt such simplification. In this paper, we resort to a widely used methods for community detection based on modularity maximization, the Louvain method (De Meo, et al., 2011). Modularity, *Q*, measures the quality of a partition of the network into communities and is defined as:

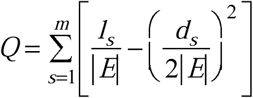

for the overall network with |*E*| edges that is partitioned into *m* communities, where *l*_*s*_ is the number of edges between the nodes belonging to the *s*-th community and *d*_*s*_ is the sum of the degrees of the nodes in the *s*-th community. The modularity maximization method detects communities by finding the network partitions that have particularly high modularities (Fig. 2A,B). Since the exhaustive search over all possible partitions is usually intractable, the Louvain Method leverages an approximate greedy optimization approach. Specifically, it iteratively optimizes local communities until the global modularity can no longer be improved, given perturbations to the current community state (De Meo, et al., 2011).

**Figure 2:**
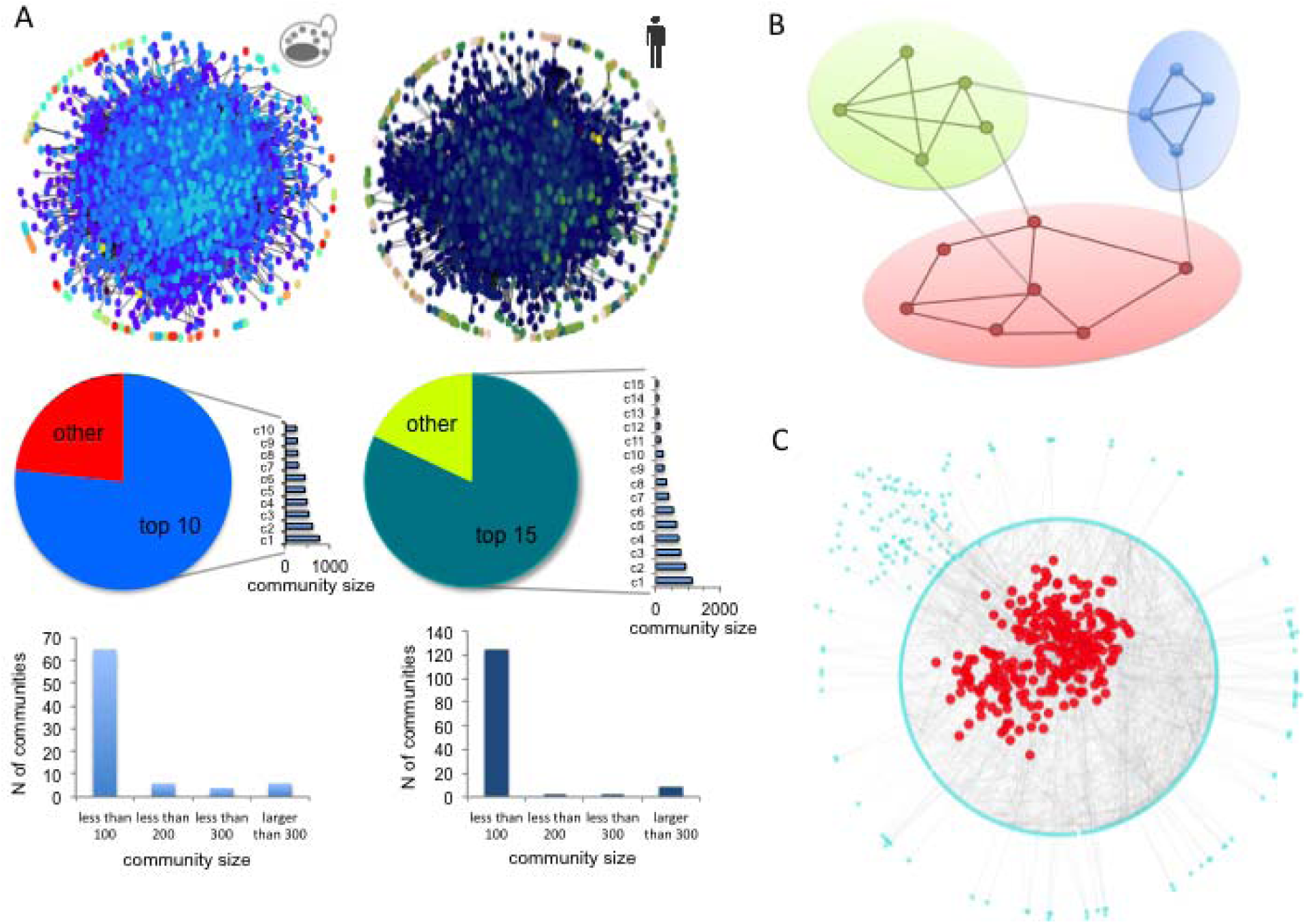
Extracting community information from protein-protein interaction (PPI) networks. A. Two PPI networks used in this works are HINT yeast network (left) and the human interactome project network (right). The networks share similarities in the size distribution of the largest communities (top 10 largest communities in yeast PPI network and top 15 communities in human PPI network, respectively, shown in the two pie charts). Furthermore, in both networks, communities with small numbers of nodes (<100) are predominant ones. B. An illustration of the basic principle behind community detection. The community detection method searches for the network partitions that have particularly high modularities using an approximate greedy optimization approach. C. The topology of a large community (red nodes) detected in the yeast PPI network and its immediate neighborhood (blue nodes). All networks are visualized using Cytoscape.

To identify genes that share similar patterns, a similarity (or dissimilarity) measure is required. However, most of the commonly used similarity/dissimilarity measures, such as Pearson correlation coefficient or Euclidean distance, do not take the prior biological information into account. In this work, we propose that such prior information on the biological network communities could be used to adjust the distance between the two gene expression profiles, thus improving the clustering performance. The weighting idea is based on a hypothesis that if two genes are a part of the same community, they are more likely to be joined via a direct or indirect interaction and hence share the same expression pattern. To achieve that, we introduce a new metric that is weighted by the community information of a gene pair in the PPI network. Specifically, for any two genes we check if these two genes are in the same community using the results of the above community detection algorithm. If they are in the same community, their original distance will be assigned a small weight with the effect of shortening the distance. Otherwise, their distance will be assigned a large weight, with the effect of elongating the distance.

The specific strategy of assigning a weight to the distance between a pair of genes is of critical importance. To derive this strategy, we take advantage on the yeast expression dataset, whose external labels correspond to the 5 phases of cell cycles. By comparing the Adjusted Rand Indices (see subsection *Evaluation criteria for two expression datasets* below for more details), one can systematically evaluate a spectrum of strategies with various magnitudes of the weights. Here, we evaluate 5×5=25 combinations of the following pairs of weights (*w*_*k*_, *w*_*m*_). The distance between a pair of genes is assigned a weight with one of the five values, *w*_*k*_=0.6, 0.7, 0.8, 0.9, or 1.0, if the genes are in the same community, and a weight with one of the five values, *w*_*m*_=1.0, 1.1, 1.2, 1.3, or 1.4, if the genes are not in the same community. The best performing weight combination will be integrated into our clustering approach.

### Deep architecture to regenerate gene expression profile

Our deep learning approach to microarray data clustering is driven by its ability to learn a hierarchical representation of the data through multiple layers of abstraction. In this work, we propose to apply our newly developed Robust autoencoder method (Zhou, 2017). The method improves the basic deep learning autoencoder model by building an outlier filter on top of a standard autoencoder, an idea that was inspired by the Robust Principal Component Analysis (RPCA) (Wright, et al., 2009).

An autoencoder is a feed-forward multi-layer neural network in which the output target is the input itself (Bengio, 2009). The method is trained to copy an input to its output. This process seems trivial, but the meaningful part is the low-dimensional hidden layers that are trained to reproduce the input. Specifically, the hidden layer is trained to be the lowest loss representation of the input constrained to some low dimensional representation. The autoencoder is a generalized framework for the non-linear dimension reduction process that is carried out by applying a non-linear activating function in the encoder and decoder parts of the method. As a result, the autoencoder can project data to a low dimensional non-linear manifold in a high dimensional space.

In contrast to the autoencoder, PCA is a basic orthogonal linear transformation method (Jolliffe, 1986). It transforms the data to a new coordinate system preserving the greatest variance. PCA projects data onto a linear subspace in a high dimensional space. However, this approach is not ideal for discovering non-linear representations, and the complexity and variability of many real-world problems often require the non-linear methods. Furthermore, in many real-world clustering tasks, the data contain outliers. Unfortunately, PCA does not work well in the presence of outliers: the linear manifold of PCA tends to shift from its otherwise optimal position, trying to offset the substantial errors caused by the outliers. The distant outliers can have a similarly profound effect on the non-linear manifold learned by the autoencoder, causing a large reconstruction error for all other data points.

To simultaneously address the problems of outliers and non-linearity, we integrate the basic ideas of Robust PCA into the autoencoder model. Robust PCA (RPCA) splits a raw input matrix *X* into a low-rank matrix *L*_*0*_ and a sparse matrix *S*_*0*_: X = L_0_ + S_0_. The low-rank matrix *L*_*0*_ represents the pattern to be extracted from the data, while the sparse matrix *S*_*0*_ consists of the outliers that cannot be captured by the low-rank pattern *L*_*0*_. We constrain the rank of matrix *L*_*0*_ to be the lowest possible, while requiring for the matrix *S*_*0*_ to be as sparse as possible. Thus, *L*_*0*_ can be represented by a linear subspace, while the *S*_*0*_ corresponds to a filter that separates the distant outliers from the linear subspace. In other words, RPCA refines PCA by making it robust to the outliers.

In the Robust autoencoder approach, we introduce a filter layer before a normal autoencoder (Fig. 3). The filter layer culls out the outlying parts that are difficult to reconstruct by the autoencoder. Thus, the outlier filter introduces robustness, while the autoencoder provides nonlinearity. The low dimensional representation learned by the autoencoder is defined by the compressed features that reflect the trend of the observation majority. Similar to Robust PCA, we decompose our input data X into two parts: *X* = *L*_*D*_ + *S*, where *L*_*D*_ is a matrix that can be represented by a non-linear manifold, and S represents the outliers which will corrupt and skew the non-linear manifold. By peeling off the outliers from *X* into *S*, the autoencoder could perfectly recover the remaining *L*_*D*_. Our loss function for a given input *X* is defined as:

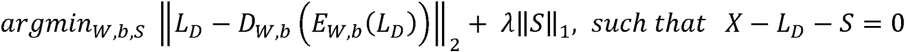

where *E*_*w,b*_ denotes an encoder function, *D*_*w,b*_ denotes a decoder function, *W* is a projection matrix, *b* is the bias term, and λ is a balancing parameter to tuning the power of sparsity. We feed *L*_*D*_ as the input data to a standard deep autoencoder to learn the low-dimensional representations. The autoencoder is trained through minimizing the reconstruction error ||*L*_*D*_ - *D*_*w,b*_ (*E*_*w,b*_(*L*_*D*_))||. The minimized reconstruction error indicates that *L*_*D*_ can be projected to a low-dimensional nonlinear manifold without significant information loss. *S* contains all outlying observations, which have high reconstruction errors and cannot be interpreted by the majority observations. We require *S* to be sparse because we want the autoencoder to capture the trend of the majority of observations, while the outliers are expected to be rare. When minimizing the first term, we want the input of the autoencoder *L*_*D*_ to be perfectly reconstructed. Thus, we need to move more observations to *S*. Similarly, when minimizing the second term, *S* will contain the increasingly smaller number of the non-zero elements. Sparsifying the outlier filter *S* leaves more errors to *L*_*D*_, and the reconstruction task of autoencoder becomes harder. In this optimization, *L*_*D*_ and *S* are mutually influenced by the constraint. The is the tuning parameter, which balances the impact of two optimizers. After training the whole model, the matrix *S* contains point-wise outliers, and *L*_*D*_ should retain the majority of information about *X* inside the hidden layer.

**Figure 3:**
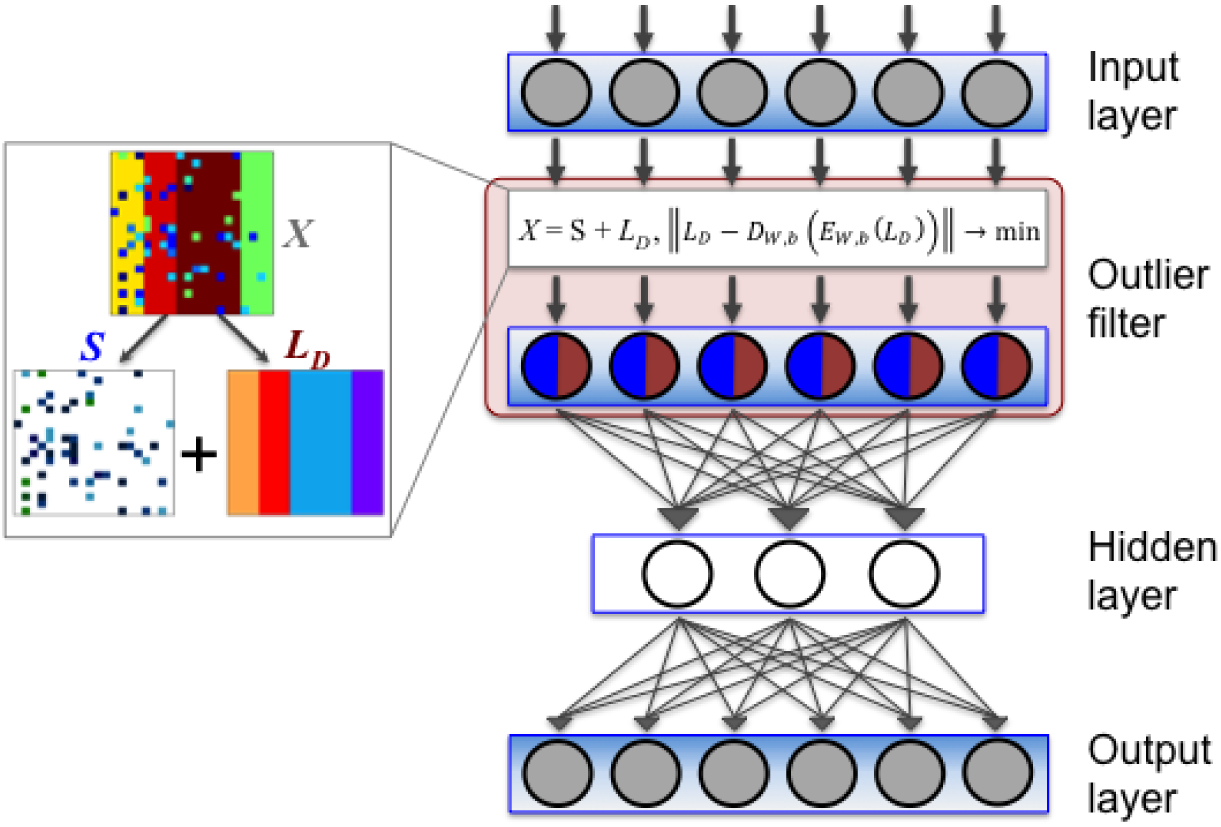
Architecture of robust autoencoder. In Robust autoencoder approach, an outlier filter layer before a normal autoencoder is introduced, providing robustness, while the autoencoder provides nonlinearity. We decompose the input data *X* into two parts: *LD*, a matrix representing by a non-linear manifold, and *S*, a matrix representing the outliers which will corrupt and skew the non-linear manifold. The goal is to filter out the outliers from *X*, thus recovering *LD*.

We solve the minimization problem of Robust autoencoder using an similar to (Zhou, 2017). While individual optimization techniques exist for training an autoencoder or Robust PCA (*e.g*., alternating direction method of multipliers, ADMM algorithm (Boyd, et al., 2011)), to the best of our knowledge no methods previously existed that could simultaneously optimize both. In (Zhou, 2017), the authors train the autoencoder using back-propagation and the outlier filter using the shrinkage function. Back-propagation is an essential element of the deep autoencoder training, but it requires the objective function to be smooth to take advantage of chain rule of differentiation. This is not the case in our problem, since the second term in our objective function, ||S||_1_, is not smooth or differentiable. However, in (Zhou, 2017) they solved this problem using a refined method is based on the basic idea of ADMM algorithm. The original objective function is broken into two smaller pieces, each of which is then easier to handle, where (1) a back-propagation algorithm is used to minimize the reconstruction cost of an autoencoder ||*L*_*D*_ - *D*_*w,b*_ (*E*_*w,b*_(*L*_*D*_))||, and (2) a shrinkage function on ||S||_1_ is used to sparsify *S* with the fixed *L*_*D*_. Then (Zhou, 2017) borrow an idea from the alternating projection forcing both optimizers to obey the constraint.

### Evaluating Robust autoencoder clustering

The gene expression data is first pre-processed using the standard data cleaning and normalization methods (Herrero, et al., 2003). Then, our new approach is introduced in two main steps. First, we use Robust autoencoder to initialize deep architectures. Once Robust autoencoders generalize specific properties of the gene expression profiles, the intermediate representation serves as an input for the clustering task. For clustering, instead of applying traditional similarity measures, we adopt the biological network based measure defined above. The measure is based on the Pearson correlation coefficient, which could detect both positive and negative correlations and is scale invariant on centered data. The similarity measure is then implemented for the agglomerative hierarchical clustering (Eisen, et al., 1998). The linkage criterion for the merge strategy in the agglomerative clustering procedure is the average linkage, which minimizes the average of the distances between all pairs of clusters.

The two clustering methods predominantly used for microarray clustering are the hierarchical clustering and partitioning clustering (Belacel, et al., 2006; D'Haeseleer, 2005). The hierarchical clustering algorithms provide a natural way for graphical representation of data. In the clustering process, each cluster is subdivided into sub-clusters, resulting in a dendrogram, in which each branch forms a group of genes sharing the similar pattern. A popular hierarchical clustering method was applied to analyze the first yeast gene expression data by Eisen *et al* (Eisen, et al., 1998); hence it is often referred as ‘Eisen clustering’. Partitioning methods, also known as non-hierarchical clustering algorithms, perform a partition of genes into a typically predetermined number of clusters so that expression patterns of genes in the same cluster are more similar to each other than those ones of genes in different clusters. k-means has been among the most popular partitioning methods (Zeger and Edelstein, 1989). The algorithm partitions around centroids, and each gene is assigned to the closest centroid. The method iterates until no genes change their cluster identity.

In the past decade, hundreds of new clustering algorithms have been developed and applied to the gene expression data. However, the performance of each clustering algorithm relies on specific properties of the input dataset and their underlying assumptions. There is no agreement on the best performing clustering algorithm for all datasets (Quackenbush, 2001). Therefore, for the baseline methods, we only implement two most widely used clustering methods: Eisen clustering and k-means. In addition, we compared our new approach to a basic autoencoder based clustering similar to the one that have been recently used for clustering the microarray gene expression data (Gupta, et al., 2015). By comparing the performance of our approach to these methods we test how much of improvement over the traditional clustering algorithms, if any, can an advanced clustering method achieve, and whether including prior biological information into the gene expression clustering analysis can further improve the clustering accuracy.

### Evaluation criteria for two expression datasets

First, we evaluate the clustering results against the reference partition for the yeast dataset, since the external labels for each gene are provided. Specifically, we use the Adjusted Rand Index (ARI) (Yeung and Ruzzo, 2001), a frequently used measure for cluster validation (Yeung and Ruzzo, 2001). ARI quantifies the degree of agreement between two partitions: one given by the clustering algorithm and the other labeled by external criteria. For a partition *U* generated by the clustering algorithm and a reference partition *V*, ARI is calculated as:

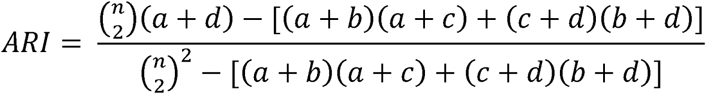

Here, *n* is the total number of samples; *a* is the number of gene pairs in the same cluster for both sets *U* and *V*; *b* is the number of gene pairs in the same cluster in *U,* but in different clusters in *V*; *c* is the number of gene pairs in the same cluster in *V* and in different clusters in *U*; and *d* is the number of gene pairs that are placed in different clusters for both, *U* and *V*. The value of ARI is defined to lie between 0 and 1, and a high score represents a good agreement between the clustering result and the reference partition. We computed the ARI scores for the clustering results using our protocol, and compared them with ARI scores obtained using the two baseline clustering methods and the basic autoencoder based clustering.

In contrast to the yeast set, no external labels are given for the GDSC sets, and the ARI metric cannot be used. In this case, a different evaluation procedure is required. Thus, we evaluate the clustering results based on their agreement with the available biological knowledge, such as Gene Ontology (Ashburner, et al., 2000). Here, we apply the following evaluation protocol. First, for the GDSC dataset, we set the number of clusters to be 100. Next, since the baseline hierarchical clustering can result in many singleton clusters, we select 10 most populated clusters for the analysis. For each cluster, we perform gene enrichment analysis and obtain the corresponding list of enriched GO terms. In the GO enrichment analysis, we use the third level of the GO hierarchy and kept the GO terms with P-value ≤ 0.01. The third level represents a trade-off between having too general, but well-populated GO terms from the second level (*e.g*., GO:0050789 regulation of biological process) and more specific but not well-populated terms from the fourth level, which cannot be used for the enrichment analysis. We compared our results for the two baseline methods. More specifically, we compared the p-values of the enriched GO terms existing for Robust autoencoder and at least one baseline method results. We expect that, for most of the significant GO terms, our protocol would output smaller p-values compared to either of the two baseline methods. These results would suggest that our protocol could identify more coherent clusters. The GO enrichment was performed using DAVID (Huang, et al., 2009), and multiple testing correction was done via false discovery rate estimation.

## Results

### Two interactomes and their corresponding community structures

Two PPI networks were extracted and analyzed, the yeast and human interactomes. For the yeast gene sets, we collected the PPI data from HINT database (Das and Yu, 2012). For the human interactome, we used the recently published interactome (referred to as HI-II-14 network (Rolland, et al., 2014)). Overall, HINT yeast network consisted of 5,687 proteins and 21,528 corresponding PPIs, while HI-II-14 network consisted of 11,787 genes and 32,465 corresponding PPIs (Table 1). A major giant component (Bollobás, 2001) existed in both interactomes, with several isolated sets of interactions on the periphery. Both interactomes shared the scale-free property (Bollobás, 2001), which means that most nodes in the network had only a few interactions and a few highly connected nodes (hubs) held the whole network together (Fig. 2A, Figs. S1, S2 in Supplementary Data).

**Table 1.** The basic statistics between the two PPI network used in the evaluation protocol.

|  | *N of genes* | *N of PPIs* | *N of communities* |
| --- | --- | --- | --- |
| HINT | 5,687 | 21,528 | 81 |
| HI-II-14 | 11,786 | 32,465 | 143 |

The detection of community structure played a critical role in our protocol. Once the interactome was constructed, we mapped the gene set to the interactome, determined which community they belonged to, and later used this information to weight the distance between any pair of gene expression profiles. We ran the Louvain method (De Meo, et al., 2011) on the two interactomes separately. After running the community detection algorithm on both networks, we obtained 81 and 143 communities from the yeast and human interactomes, correspondingly (Table 1, Fig. 2A). The largest community in the yeast interactome was composed of 764 genes. The top 10 largest communities covered 77% of the total proteins in the network. The other communities were all composed of only few nodes. Similarly to the yeast network, the first 15 communities accounted for 82% proteins in the human interactome, while the largest community contained 1,129 proteins (9.6%).

### Incorporating prior biological network information and finding a proper weighting strategy

After the community detection stage, we examined every gene pair from the gene expression list to determine if they were in the same community. Then, we utilized this information to weight the distance between each pair of gene expression profiles. We compared the weighted clustering results with the baseline clustering results to demonstrate the effectiveness of incorporating network community information. For the baseline clustering methods, we implemented two most widely used approaches, k-means and hierarchical clustering. The two baseline methods were considered as the “un-weighted” clustering approaches. We then determined the optimized combinations of weights using a basic grid search on the hierarchical clustering method. Specifically, the search explored the weights from the range 0.6 to 1 (with a step of 0.1) for each pair of genes that were in the same community, and from the range 1.0 to 1.4 (with the same step) if the genes were not in the same community. The best performing combination was selected for our protocol.

The effectiveness of including the biological information was assessed on the labeled yeast gene expression dataset, since one could accurately evaluate the clustering performance only when the external labels were available (Fig. 4A, 4C). For each of the two baseline methods, we set the number of generated clusters to be five (matching the total number of different labels in the yeast dataset). Hierarchical clustering method performed with ARI of 0.448 on the yeast dataset, while k-means performed with ARI of 0.420. The ARI values after applying different weighting strategy ranged from 0.444 to 0.488 (Fig. 4A, Table 2). Overall, the accuracy after applying the weighting strategy was better compared to the un-weighted baseline methods. These results demonstrated that the biological network community information could be utilized to improve the traditional clustering. The results also supported the hypothesis that gene pairs in the same community of the PPI network are more likely to share the same expression pattern. Also, we note that the weight combination 0.9 and 1.3 yielded the most accurate results. Therefore we adopted this weighting strategy for our protocol.

**Table 2.**
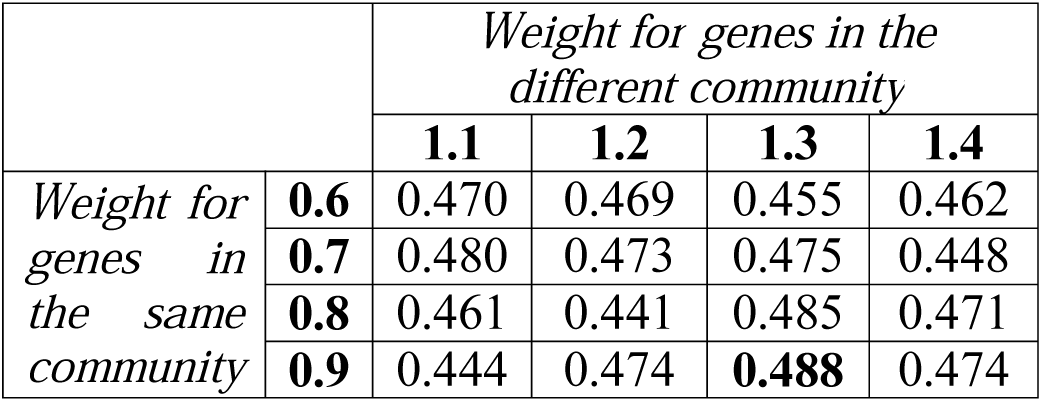
Comparison of results from different weighting strategies obtained when including the network community information.

|  |  | *Weight for genes in the different community* |  |  |  |
| --- | --- | --- | --- | --- | --- |
|  |  | **1.1** | **1.2** | **1.3** | **1.4** |
| *Weight for genes in the same community* | **0.6** | 0.470 | 0.469 | 0.455 | 0.462 |
|  | **0.7** | 0.480 | 0.473 | 0.475 | 0.448 |
|  | **0.8** | 0.461 | 0.441 | 0.485 | 0.471 |
|  | **0.9** | 0.444 | 0.474 | **0.488** | 0.474 |

**Figure 4:**
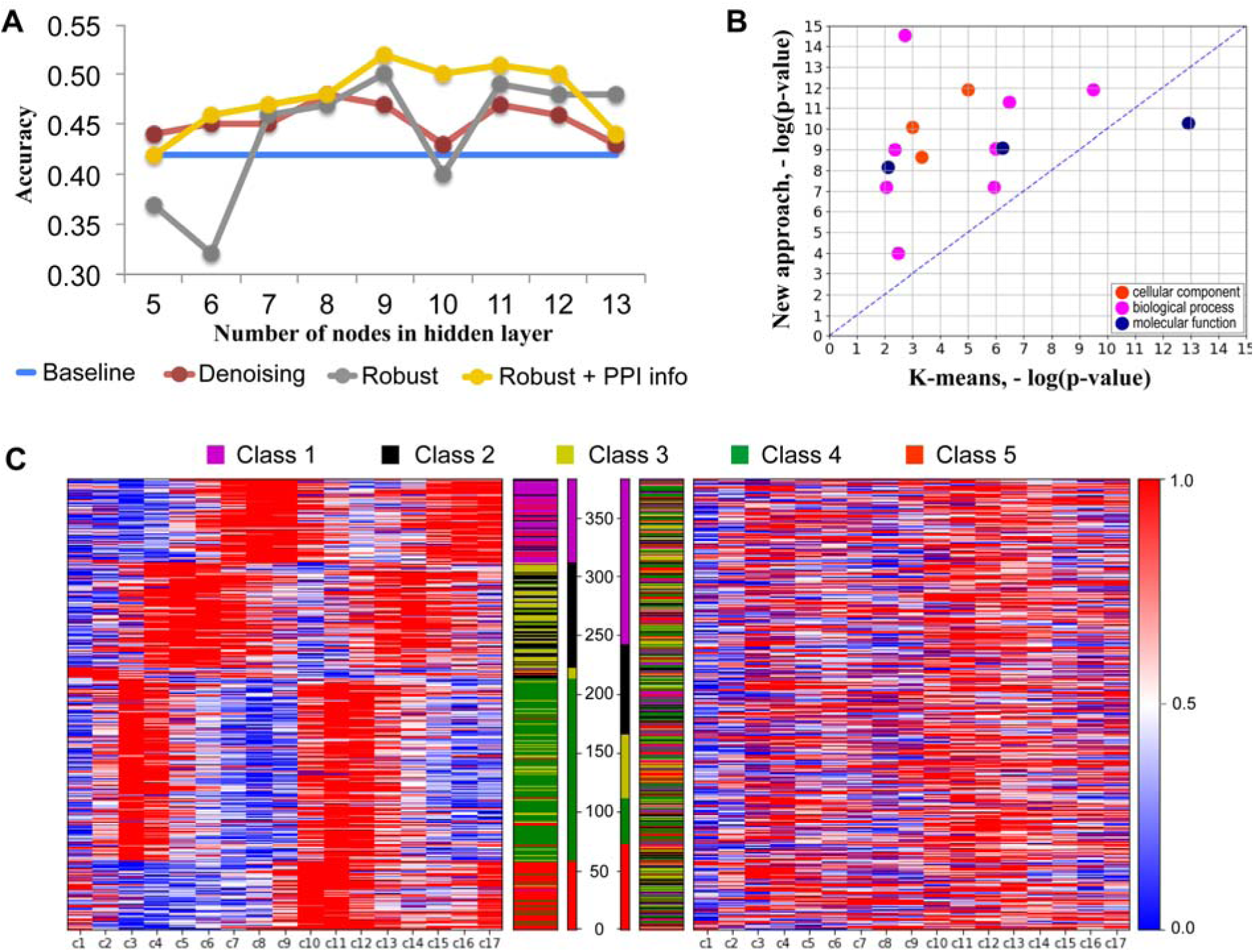
Evaluation of the new clustering approach. **A.** Comparison of the performance of two deep architectures against baseline methods performed on previously labeled yeast gene expression dataset. The accuracy measure used here is Adjusted Rand Index (ARI). Shown is the comparison of our approach that combines the Robust autoencoder architecture with the PPI network community information (yellow) against the base line K-means clustering method (blue), standard denoising autoencoder (red), and Robust autoencoder without additional biological information (grey). **B.** Comparison of enriched Gene Ontology terms between our approach and K-means for the human gene expression dataset. The values are converted using negative log of p-value function. A smaller p-value reflects a larger proportion of the cluster members sharing the same GO term. **C.** Performance of our approach (left) against the base line K-means clustering (right) on the yeast gene expression dataset provides a visibly better clustering into 5 previously labeled gene classes across 17 different time stamps (c1-c17).

The denoising autoencoder model (Vincent, et al., 2008) is another popular deep learning architecture and can be viewed as a stochastic version of the autoencoder. It randomly corrupts the input data and trains the parameters to recover the uncorrupted data from the corrupted one. Denoising autoencoders can be stacked to form a deep network, *i.e.* stacked denoising autoencoder. The denoising autoencoder’s goal is to learn the mapping from the corrupted data to the original uncorrupted data. One of the method’s caveats is that it still needs the information about the original uncorrupted data for the training. Since the original, uncorrupted, data present the crucial prior knowledge for denoising autoencoder, the quality of the original data will influence the denoising autoencoder’s map building and the quality of discovered features. If the original input contains outliers, denoising autoencoder’s training will still learn to recover these outlying parts and the quality of discovered features could be misled by these outlying parts.

In contrast, Robust autoencoder distinguishes the outliers from corrupted data without the knowledge of uncorrupted data. To illustrate that Robust autoencoder is a better choice than the denoising autoencoder for regenerating the gene expression profile, we applied both methods on the yeast expression dataset. We considered the individual effects of deep architecture on the clustering results, *i.e.*, without applying the community information to weight the distance in the protocol. For Robust autoencoder, the best ARI obtained across different hidden layer sizes was 0.5, whereas the highest ARI obtained for the denoising autoencoder was 0.48 (Fig. 4A). Thus, our deep architecture performed better, although not significantly. We also noted that Robust autoencoder suffered from the greater variation of ARI values compared to denoising autoencoder.

### Evaluation of our protocol on the yeast gene expression dataset

In our protocol, instead of taking as an input for clustering the raw expression data, we reconstructed the features via Robust autoencoder and used this intermediate feature representation for clustering, so the best performing weight combination was not directly assigned to the raw dataset. To compare the results of our protocol with the baseline methods on the yeast dataset, the same ARI measure was calculated. The results showed that our protocol, which incorporates the prior biological information on the regenerated data from the deep architecture, outperformed the baseline methods applied to the raw data (Fig. 4A, 4C, Fig. S4 in Supplementary Data). Furthermore, the results of our protocol outperform the baseline method with the used community information for the pairs of genes. This behavior is perhaps due to the ability of the architecture to learn important properties in the underlying input distribution. Also, we note that, compared against the results without incorporating biological information, the former clustering results had smaller variation of ARI values, suggesting that incorporating the prior biological information could stabilize the clustering process. Finally, we found that deep architecture does not guarantee that it will always perform better than the basic clustering methods. For instance, our deep architecture with hidden size of 5, the performance is comparable to the baseline methods. This implies that tuning parameters of deep architecture is a critical but not a simple step for these methods.

### Evaluation of our protocol on the human gene expression dataset

When implementing our protocol on the GDSC dataset, we used the results got from the Yeast dataset to guide the construction of the deep architecture. Specifically, we used a comparable percentage of the input layer size as in the best performing deep structure for the Yeast dataset to build the hidden layer. This led to a hidden layer with 55 nodes.

The human gene expression dataset consisted of 17,419 genes expressed in 83 cell lines. We independently applied our protocol as well as the k-means and hierarchical clustering methods on this gene set, while setting the cluster number in each case to be 100. Out of 100 clusters, we focused on the top 10 largest clusters and performed the GO enrichment analysis on these clusters. We only selected the third level GO terms in the GO hierarchy tree and compared the results against k-means and hierarchical clustering (Fig. 4B, Fig. S3 and Tables S1, S2 in Supplementary Data). Comparing against k-means, 22 GO terms from the third level were enriched in at least one cluster in both cases, and most of the GO terms identified by our protocol had smaller *P*-values. This indicated that our protocol could group a more coherent and meaningful set of genes into a cluster. Compared against hierarchical clustering, we obtained 114 GO terms enriched in at least one cluster. In this case, the number of GO terms obtained in our approach (*N*_1_=61) with smaller *P*-value was slightly larger than the number obtained in hierarchical clustering (*N*_1_=53). This did not indicate that our protocol could significantly improve the traditional hierarchical clustering in terms of generating more coherent clusters. However, we noted another interesting observation. One main problem about hierarchical clustering is that it groups too many genes into a very large, giant, cluster. In this case, the largest cluster resulted from hierarchical clustering consisted of 11,043 genes, and its size was almost comparable to the first three largest clusters found by our protocol. This suggests that our protocol could compensate the inability of hierarchical clustering to further separate the clusters.

## Discussion

In this paper, we proposed a novel approach to microarray-based gene expression clustering that combines a new deep learning approach with the prior biological knowledge. We trained a Robust autoencoder to learn interesting characteristics of the gene expression profiles. The obtained low dimensional representations of gene expression profiles were then used for the clustering task. To increase the clustering accuracy, the clustering algorithm employed a knowledge-based molecular network similarity measure. We compared the performance of our clustering approach with two traditional and commonly used clustering methods, k-means and agglomerative hierarchical clustering, on two distinct gene expression datasets. Our results demonstrated the effectiveness of using (i) deep networks and (ii) prior biological information for the gene expression clustering analysis.

Several conclusions have been made from this work. First, we used a fairly simple deep learning architecture because of the long computation time. In future work, we plan to adopt a much deeper architecture. An autoencoder with a single encoder and decoder is usually considered as a shallow model. The way of extending shallow autoencoder to deep autoencoders is to add more encoding and decoding phases. A typical implementation of this idea is the stacked autoencoders [42]. The same idea could be applied to the Robust autoencoder model presented here. To address the problem of computational overhead, one can resort to the GPU computing algorithms.

Second, in spite of the improved accuracy over the standard clustering methods as well as over the basic autoencoder, our clustering protocol could be further optimized in several ways. For example, one can explore other distance metrics that have been previously shown to perform well in the clustering with homogenous features [5]. Alternatively, we plan to investigate if the clustering performance can be improved by supplying the complementary biological information. For example, instead of the gene community information used in this work, the shortest path between two nodes in the network can be considered, since the former sometimes provides more accurate information than the latter.

## Acknowledgements

This work is supported by National Science Foundation (DBI-1458267 to DK).

## Author Contributions Statement

HC conceived the idea. HC and DK designed experiments. CZ and RP developed Robust Autoencoder. HC, XD and YL collected data and performed the experiments. HC and DK analyzed the results. RP supervised the work of CZ. DK directed the study and supervised the work of HC. HC and DK wrote the manuscript, with all authors contributing to discussion and revision of the drafts.

